# Cre-Lox miRNA-delivery technology optimized for inducible microRNA and gene-silencing studies in zebrafish

**DOI:** 10.1101/2024.02.22.581357

**Authors:** Fangfei Guo, Alisha Tromp, Haitao Wang, Thomas E Hall, Jean Giacomotto

**Author notes:** Contributed equally to this work (Fangfei Guo, Alisha Tromp).

## Abstract

While many genetic tools exist for zebrafish, this animal model is still lacking a robust gene-silencing and microRNA-delivery technology enabling spatio-temporal control and reliable traceability. We have recently demonstrated that the use of engineered pri-miR backbones can be used to both trigger stable and heritable gene knockdown and/or express microRNA(s) of choice in this organism. This approach/material fills a gap that exists in the zebrafish genetic toolbox to study microRNA biology and has the potential to be a powerful complementary tool to CRISPR-knockout and CRISPR-interference approaches by enabling conditional, traceable tissue-specific and polygenic gene-silencing. However, to date, this miRNA-expressing technology still presents important limitations. First, to trigger potent knockdown(s), multiple synthetic-miRNAs must be expressed simultaneously which compromises the co-expression of fluorescent marker(s) and knockdown traceability, thereby reducing the interest and versatility of this approach. Second, when the gene knockdown triggers significant phenotypes, like homozygous mutants with severe early phenotypes, it is difficult, if not impossible, to maintain adult transgenic carriers. Here, to solve these problems and provide a mature RNAi and microRNA-delivery technology for zebrafish research, we have generated and validated new RNAi reagents and an inducible delivery system based on the Cre/Lox technology. This new system allows the creation of asymptomatic/silent carriers, easing the production of embryos with potent knockdowns that can be traced and spatiotemporally controlled. Following this technological development, we demonstrate the utility of our approach by generating novel inducible models of spinal muscular atrophy (SMA) that open new avenues for studying the normal and pathogenic functions of the smn1 gene, as well as for establishing large-scale screens. Finally, and very importantly, the materials and techniques developed here will simplify research into microRNA biology, as well as conditions involving haploinsufficiency and multiple genes.

## INTRODUCTION

The zebrafish has become a powerful animal model for investigating gene function, modelling human diseases and screening for bioactive compounds. It combines the tractability and versatility of *C. elegans* and *D. melanogaster* with the ability to replicate/model human disorders and enabling screens for drug discovery and chemical genetics. Nonetheless, one key technology which has been missing and remained intractable is a robust methodology for RNA interference (RNAi, gene-silencing or knockdown) and microRNA-delivery^1-4^. To develop such tools for zebrafish, we have recently tested a variety of approaches and found that engineered *pre-miR* can be used to express any mature miRNA(s) of choice for either microRNA basic research studies and/or to silence endogenous genes-of-interest throughout the entire lifespan of the fish^5-7^. In practice, and as presented previously^5^, a RNAi cassette containing i) a polymerase-II promoter, ii) a fluorescent marker and iii) a custom *pri-miR*(s) (designed to release a synthetic microRNA (miR) against the 3’UTR of a gene-of-interest) is cloned into a tol2-transgene for genomic integration^8^. Upon activation, this transgene triggers co-expression of a fluorescent marker and the synthetic-miR(s), thereby silencing the targeted gene(s) while highlighting/marking the affected cells **(fig. S01A)**. Although this technology opens new avenues of research for zebrafish genetics, unfortunately, to date, this RNAi technology still presents important limitations to apply to any gene-of-interest and to multi-genic knockdown. <u>First</u>, to trigger potent knockdown, one must express multiple synthetic miRNAs against different target sites located on the gene-of-interest’s 3’UTR. To achieve this, multiple *pri-miRs* must be chained as concatemer **(fig. S02A)**. However, after more than 3x chained *pri-miRs*, we found it difficult to generate transgenic lines with visible co-expression of fluorescence (see **Table 01** below as an example for lines targeting the gene *smn1*). This loss of fluorescence is likely due to the requirement for the *pri-miRs* to first be processed/cut via Drosha and Dicer to generate mature miRs. These successive cuts leave the associated mRNA without polyA tails thereby triggering their denaturation (**fig. S01A and fig. 01B**). In this model, the miRs processing/maturation directly competes with the translation of the attached fluorescent marker, with the more *pri-miRs*/miRNAs chained, the less fluorescent protein successfully produced. <u>Second</u>, when the gene knockdown triggers significant phenotypes, similar to homozygous mutants with strong early defects, it becomes challenging to maintain stable transgenic lines with even one copy of the RNAi-transgene. This can be a significant problem when one tries to model human diseases such as Spinal Muscular Atrophy (SMA)^5,6,9^ which are associated with SMN haploinsufficiency/partial-LOF. Expression of miRNAs against the causal gene *smn1* triggers strong early motor neuron defects, motor function loss and premature death consistent with disease presentation. However, the more severe the phenotype, the more difficult it becomes to both isolate and maintain animals.

**Table 1:** Generation of conditional smn1-LOF animals based on previous RNAi cassettes incorporating an intronic sequence or not. This table presents the number of animals screened/selected post-injection as well as the number of founders identified. All the founders have been compared and ranked based on the BFP expression/brightness of their progeny, with a score of 10 for the brightest. Only the six brightest lines were conserved and outcrossed with tg(UBI:iCre) to evaluate the conditional system efficiency and the associated red fluorescence.

| Responder lines generation | Number of selected F0 embryos | Number of F0 adult | Number of Positive F0 founders | Selected F1 lines | Blue brightness (unfloxed) | Red brightness (floxed) |
| --- | --- | --- | --- | --- | --- | --- |
| tg(UBI:loxBPF:dsRED:miRsmn1-4141) | 50 | 38 | 14 | #1 | 10 | 0 |
|  |  |  |  | #2 | 7 | 0 |
|  |  |  |  | #3 | 6 | 0 |
|  |  |  |  | #4 | 6 | 0 |
|  |  |  |  | #5 | 6 | 0 |
|  |  |  |  | #6 | 6 | 0 |
| tg(UBI:loxBPF:mKate:INTRON_miRsmn1-4141) | 85 | 52 | 11 | #1 or tg(floxSMN) | 10 | 10 |
|  |  |  |  | #2 | 7 | 9 (Mosaic) |
|  |  |  |  | #3 | 7 | 8 |
|  |  |  |  | #4 | 6 | 7 |
|  |  |  |  | #5 | 6 | 5 |
|  |  |  |  | #6 | 4 | 6 |

**Figure 1:**
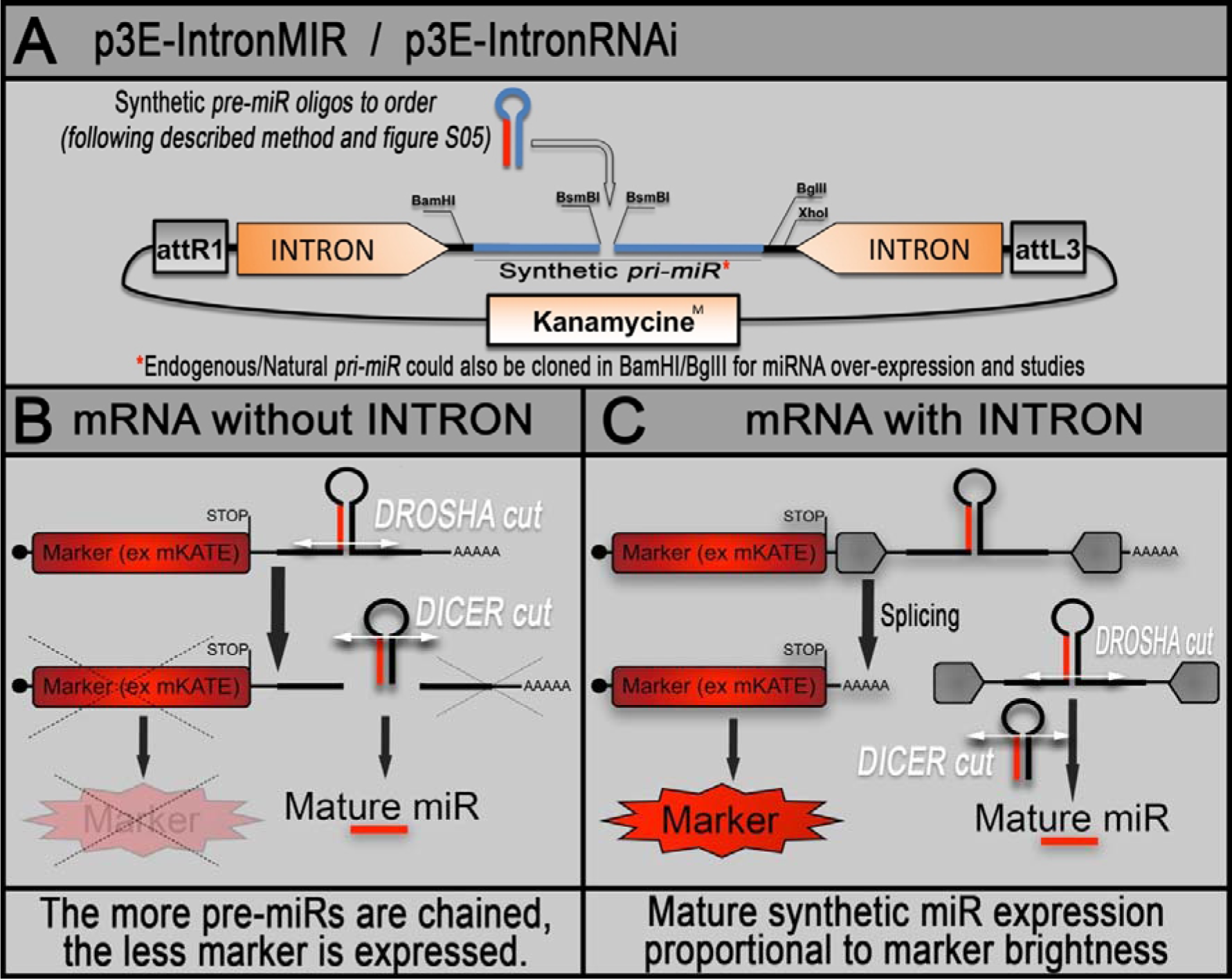
Schematics of the new RNAi backbone for zebrafish gene silencing. **A,** Tol2kit compatible p3E presenting an optimised empty synthetic *pri-miR* (or RNAi cassette) embedded into a β-globin intronic sequence. This presented *pri-miR* is designed to allow rapid directional insertion of synthetic *pre-miR* of choice. Synthetic p*re-miR(s)* of choice are generated by annealing specific top and bottom RNAi oligos designed to release a mature miRNA directed against the 3’UTR of a gene-of-interest (see method section and previous material^5^). The RNAi cassette is flanked by restriction sites allowing subsequent and repetitive chaining (see method and **fig. S2**) for generating p3E-RNAi plasmid with multiple *pri-miR* for either increasing the potency of the desired knockdown or targeting multiple genes at the same time. **B,** Without intron, the *pre-miR*/RNAi cassette is transcribed along with a co-marker on the same RNA, which is cut by Drosha in the nucleus for releasing the associated *pre-miR*. This cut leaves the mRNA without polyA tail and leads to its rapid degradation. The associated fluorescence is not a good indicator of the activity and amount of synthetic miRNA produced. **C,** The presence of the intronic sequence is supposed to help rescue the co-expression of the marker (**see also figure S1**).

To solve these problems and deliver a mature RNAi and microRNA-delivery technology for zebrafish, we have generated and validated i) new reagents that are compatible with the zebrafish tol2kit and that offer the possibility to chain numerous synthetic or natural *pre-miR* without affecting the co-expression of a fluorescent marker as well as ii) an inducible system based on the Cre/Lox technology allowing the generation of conditional lines that can be induced with spatiotemporal control. Subsequently, using this system, we generated a versatile conditional zebrafish model of SMA. This model is opening new avenues for investigating SMN function and for large-scale experiments such as drug screenings.

Finally, this system promises to be very useful to the community to express and study endogenous miRNA(s).

## RESULTS

### Generation of an optimised kit for both RNAi and endogenous/natural miRNA expression

We first sought to develop a technology where strong and robust expression of our selectable fluorescent marker is able to occur together with multiple concatenated *pri-miRs*. We hypothesized that the RNAi expression cassette could be incorporated into an intronic sequence, leading to its splicing during mRNA maturation. This would result in a mature mRNA encoding the fluorescent marker terminated by a polyA tail alongside the spliced RNAi cassette(s) (**fig. 01 and fig. S01**). Supporting this rationale, nearly half of the known human endogenous miRNAs are encoded within the introns of protein-coding genes^10^. We reviewed the literature to find an appropriate sequence to use and selected the rabbit β-globin intron that has been demonstrated multiple times to be efficiently spliced in zebrafish while, at the same time, providing a significant increase in protein translation/expression^11-13^. We assembled this sequence followed by an SV40 polyA signal within a tol2kit empty p3E-vector carrying a kanamycin resistance cassette. Since the kanamycin resistance gene possesses a BsmBI restriction site that is incompatible with RNAi cassette assembly, we mutated this site so the final plasmid, named p3E-IntronRNAi (Addgene #163380, also called p3E-IntronMIR), would present only the two BsmBI sites required to introduce the synthetic *pre-miRs* (**fig 1A**, and method section)^5^. Note that the RNAi cassette is surrounded by unique restriction sites (BamHI-BglII/XhoI) enabling straightforward and endless chaining (**fig S02**).

Finally, to enable the expression and the study of endogenous miRNA, one can easily use the present BamHI/BglII sites to swap the present RNAi cassette with an amplified human or zebrafish *pri-miRNA*, surrounded by similar sites for keeping chaining versatility (**fig. 01**). On can either order these *pri-miRNA* as gene-blocks or amplify them from the respective genomes using compatibles oligos.

### Validation of the intron-based miR-delivery and RNAi approach

To both test that i) the miR/RNAi-delivery cassette is still functional when placed within an intron and ii) that several *pri-miR* could now be chained without compromising the stability/co-expression of a fluorescent marker, we <u>first</u> conducted transient experiments. We took advantage of a transgenic line already available in the laboratory, *i.e.* tg(UBI:mCherry:SP137), which ubiquitously expresses an mRNA encoding mCherry and a custom 3’UTR acting as a sensor for endogenous and/or synthetic miR-137^14^. This line presents strong and homogeneous ubiquitous red fluorescent expression. We designed a synthetic *pri-miR* directed against this transgene 3’UTR. As presented in Figure 01 and in the method section, we first built a single-repeat 1x-RNAi cassette by inserting a custom *pre-miR137* into both the miR-delivery plasmids with or without intron. Following a traditional 4 component gateway LR reaction (**fig. S02B**), we then generated two tol2 1x-RNAi-transgenes encoding an eGFP and RNAi cassettes (with or without intron) under the control of a striated muscle-specific promoter (503UNC)^6^, respectively named 503UNC:eGFP_RNAi_1x_miR137 (or 153) and 503UNC:eGFP_RNAi_INTRON_1x_miR137 (or 155). A control transgene has also been generated using an RNAi cassette encoding non-targeting miRs. Those plasmids were injected along with transposase into tg(UBI:mCherry:SP137) as presented in **figure S03**. If functional, excepting for the control, the injected transgenes should lead to the co-expression of both eGFP and the designed synthetic-miR137 directed against the integrated UBI:mCherry:SP137 transgene, silencing its expression cell-specifically in striated muscle cells. At 4day-post-fertilisation (dpf), we observed that all transgenes led to detectable GFP expression evidenced by mosaic green fluorescence, as expected for a transient DNA injection (**fig. 02**). All transgenes but the control also led to mosaic loss of mCherry expression, with a negative correlation/overlap with eGFP expression (**fig 02D, 02H, 02L**), demonstrating proper cell-specific silencing of the targeted transgene and, thereby, validating that the presence of the intronic sequence did not interference with the maturation of the designed synthetic miRNAs (**fig. 02**). <u>Second</u>, we tested if the intron would help improving/rescuing the stability of the co-marker expression upon chaining several *pri-miR*/RNAi cassettes. Following a similar cloning strategy but chaining 4x *pri-miR137* together, we generated two new transgenes named 503UNC:eGFP_RNAi_4x_miR137 (or 154) and 503UNC:eGFP_RNAi_INTRON_4x_miR137 (or 156) (**fig. S02**). As observed for the 1x-constructs, both plasmids successfully triggered eGFP expression and mosaic decrease/extinction of red fluorescence, demonstrating proper transcription of the transgenes and proper maturation of the associated *miR137* miRNAs (**fig. 03**). However, it appeared obvious that the 4x-*pri-miR* construct without intron triggered loss of red fluorescence that was not associated with eGFP expression. This observation was quantitatively confirmed as presented in **figure 03I**, *i.e.* decrease of red fluorescence did not negatively correlate with green fluorescence emission, evidencing that eGFP expression can be lost when several *pri-miRs* are chained. In contrast, when the 4x chained *pri-miRs* were embedded into the β-globin intron, one could still observe a negative correlation between loss/decrease of red fluorescence and presence/increase of green fluorescence, demonstrating that i) the *pri-miR* repetitions did not interfere anymore with the expression of the marker cassette (here eGFP) and ii) the presence/intensity of the fluorescent marker reflects the level of miR-expression. These conclusions were also supported by the transgenic/stable experiments presented below (**table 1**).

**Figure 2.**
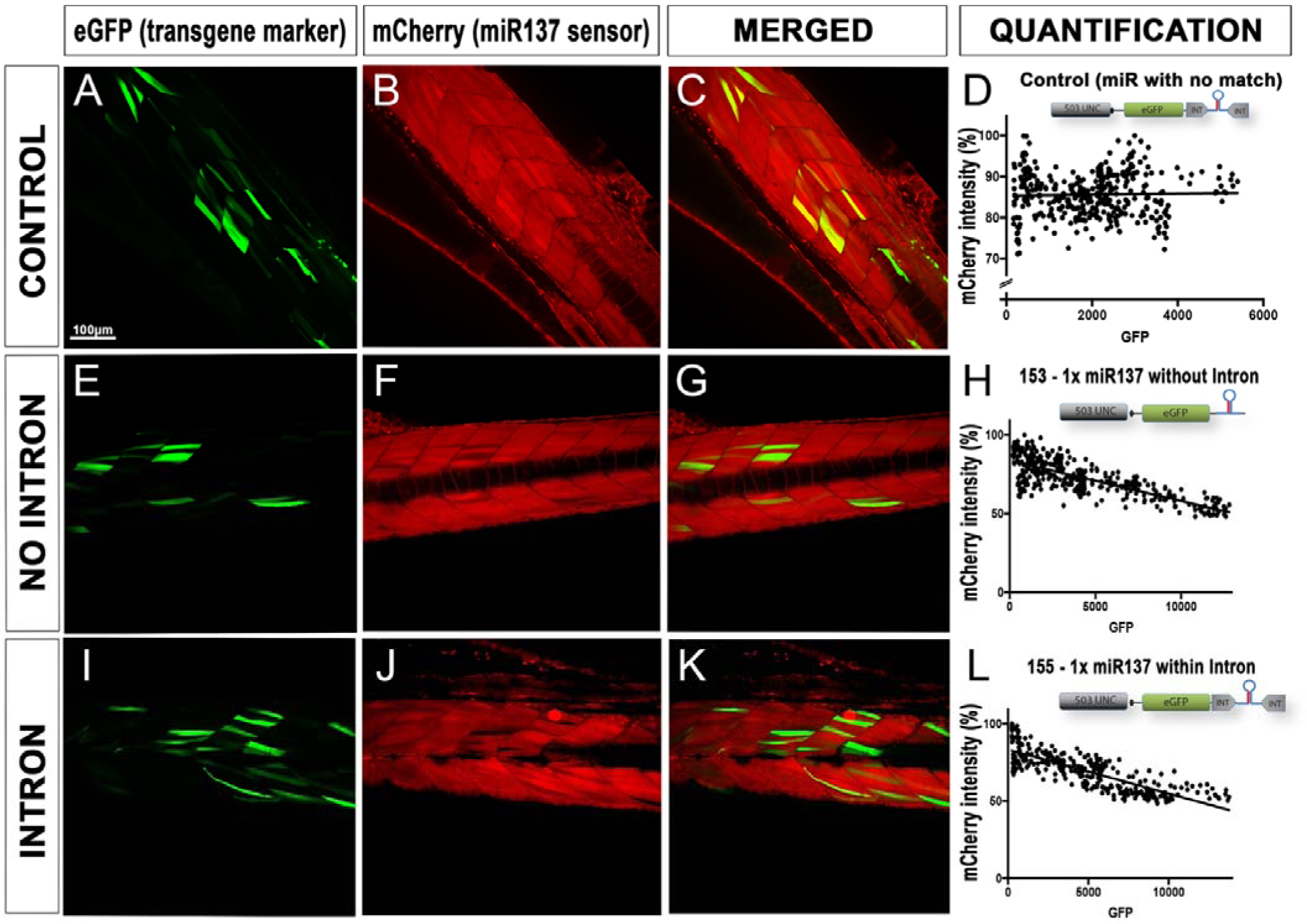
Validating effect of intron based RNAi approach on endogenous miRNA processing. **A-C, E-G, I-K** Ubiquitously expressing mCherry fluorescent embryos from Tg(UBI:mCherry:SPmiR137) cross were injected with tol2 transposase and 50pg of anti-miR137 expressing constructs - 153 (without intron), 155 (with intron) and 157-control. All injected plasmids led to mosaic eGFP expression. Plasmids 153 and 155 (not 157-control) led to knockdown of mCherry sensor protein. A minimum of 6 larvae were analysed per condition to estimate knockdown efficiency. Graphs **D, H** and **L** are representations of pixel intensities (each dot presents a grey value) of green versus red fluorescence (presented as a percentage) from z-stacks imaged with a confocal microscope at 4dpf. Scale bar is 100µm.

**Figure 3.**
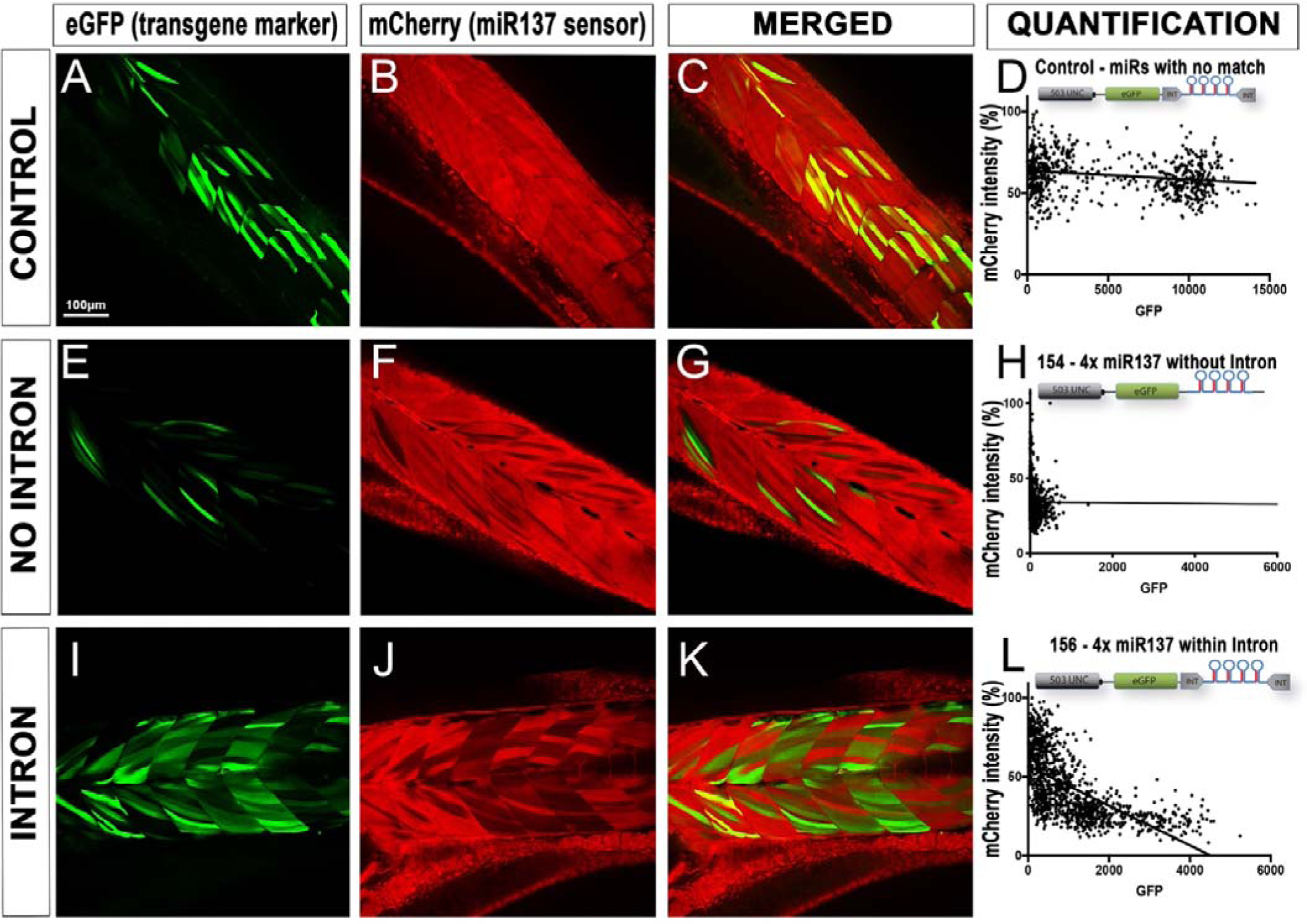
Validation of intron based RNAi approach on rescuing co-expression of fluorescent markers. **A-C, E-G, I-K** Heterozygous Tg(UBI:mCherry:LmiR137) embryos with ubiquitous mCherry expression were injected with tol2 transposase and 50pg of 4x anti-miR137 miRNAs construct without intron (plasmid 154) and with intron (plasmid 156), including a control (plasmid 157). A minimum of 5 larvae were imaged at 4dpf using a confocal microscope to correlate the presence of 4x anti-miR137 expressing eGFP (with and without intron) to decrease/absence of mCherry fluorescence. Pixel intensities (gray values) of green versus red fluorescence (presented as a percentage) plotted in graphs **H** demonstrate no correlation of fluorescence in embryos injected with plasmid 154, similar to control 157. Comparatively, a linear correlation of green versus red fluorescence **(L)** is evidenced in embryos injected with plasmid with intron (plasmid 156).

### Generation of a robust inducible miRNA-expressing and RNAi system

Following the optimisation and validation of the new miR-delivery/RNAi cassette, we worked on establishing a robust inducible system. We previously used the Gal4/UAS technology but experienced important drawbacks that strongly hampered the flexibility and potential of this system^6^. We experienced i) random and strong methylation-induced silencing of the responder/miR-delivery transgenes, generating heterogeneous results difficult to analyse, and ii) toxicity problems with ubiquitous expression of Gal4. That being said, one advantage of the Gal4/UAS system was its ability to trigger strong expression of the responder/miR-delivery cassettes, thereby limiting the need to use long concatemers to achieve potent knockdown^6^. Nonetheless, the problem of ubiquitous expression combined with random silencing generated too great limitations for both our basic research and our drug discovery programs. We thereby worked at establishing a more reliable conditional system for which we introduced the Cre/lox system^15^.

This alternative inducible genetic system is also based on driver and responder transgenes (**fig. 04**). The driver transgene/line is used to express Cre-recombinase in the desired tissue(s) and/or at the desired time-points in order to trigger Lox-recombination of the responder transgene(s); leading to either sequence deletion or inversion depending on the Lox-design/orientation. To establish the driver transgenes and lines used in this study, instead of using the Cre-recombinase commonly used in the zebrafish community, we incorporated the codon-improved Cre (iCre) into a tol2-kit compatible pME (*i.e.* pME-iCre, addgene #171792)^16^.

**Figure 4:**
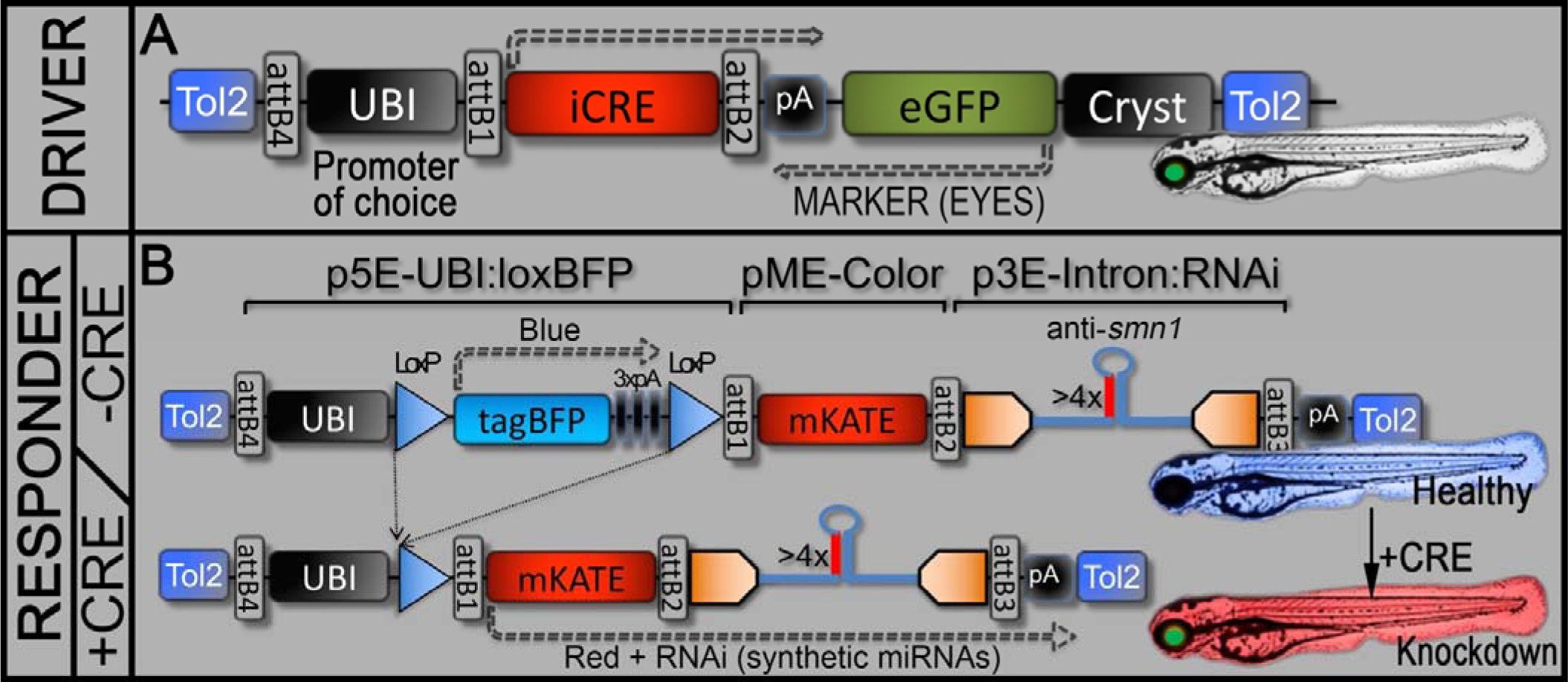
Conditional Cre/Lox RNAi genetic system. **A,** Schematic representation of the driver transgene, with here a ubiquitous promoter, named UBI:iCre. Tissue-specific promoter can obviously be used. **B,** Schematic representation of the silent/conditional gene-silencing responder transgene. In the absence of iCRE, only BFP will be expressed and the animal will remain unaffected by the RNAi cassette, thereby could be maintained without special care. In the presence of iCre, genetic recombination will lead to excision of the BFP cassette and expression of mKate along with the miRNAs of choice; with here 4x anti-*smn1* miRNAs, resulting in potent zebrafish *smn1* knockdown. This recombination is irreversible.

We further built destination clones with ubiquitous, muscle, motoneuron or pan-neuronal promoter (Ubiquitin, 503UNC, MN, or HuC promoter respectively^6,17,18^), along with a marker cassette encoding eGFP under the control of a crystallin promoter (Green expression in lens/eyes for identifying transgenic carriers) (**fig. 04A**). This iCre version has been previously optimised for improving expression, stability, and translocation to the nucleus^16,19^. The corresponding driver transgenic lines have been generated through a traditional tol2-transgenesis process, as described in the method section. Animals were selected and maintained thanks to eGFP expression in their eyes/lens.

To establish the conditional RNAi-responder transgenes, taking inspiration from the (Ubi:Switch) system used for cell lineage^20^, we built a genetic system allowing the integration of the miR-delivery/RNAi cassette in a silent state, and activable upon expression/presence of iCre (**figure 04B**). For that, we built a p5E:UBI:lox:tagBFP:lox (see method section) that can be recombined with any additional color/marker (mKate here in our study; Red fluorescence) and any miR-delivery/RNAi cassette of choice. In practice, this system allows to generate asymptomatic/silent-RNAi responder carriers expressing BFP ubiquitously and that can be activated upon expression of Cre/iCre. In the presence of Cre/iCre, the BFP cassette would be floxed/excised triggering expression of mKate along with the miR-delivery/RNAi cassette, meaning the animals would turn from healthy/unaffected blue fluorescent to knockdown red fluorescent larvae. Interestingly, this new system eases the selection of lines with strong expression of the RNAi cassette without having to deal with negative selection pressure (due to the potential phenotypic effect of the associated knockdown). Indeed, one can now easily screen animals based on BFP expression (blue fluorescence) to select carriers with strong transcriptional activity.

### Proof-of-principle and generation of a versatile model of SMA

To validate the robustness of our inducible system, we further tried to establish a conditional model of SMA. We previously validated an anti-*smn1* RNAi cassette presenting 4x *pri-miR* repeats (named miRsmn1-4141)^5^. We used that material to build two anti-*smn1* responder constructs/transgene as presented in **figure 4B**. Both constructs are similar but one present the 4x RNAi block embedded into the aforementioned p3E-IntronRNAi. We named those anti-*smn1* transgenes UBI:loxBFP:dsRED:miRsmn1-4141 and UBI:loxBFP:mKate:INTRON_miRsmn1-4141. Both were injected at one cell-stage of tg(MN:GFP) embryos (transgenic line expressing eGFP in motor neurons)^17^ along with transposase mRNA for random genomic integration. Fifty F0 embryos were screened based on visible tagBFP expression and raised until adult stage. Adults F0 were outcrossed against wild type (wt) in order to identify founders that successfully transmit the injected transgene. Fourteen (14) founders out of 38 F0 adult screened adults were identified for UBI:loxBFP:dsRED:miRsmn1-4141 (no intron) and 11 out of 52 for UBI:loxBFP:mKate:INTRON_miRsmn1-4141 (intron) (**table 1**). We further compared and ranked those founders on a scale of 1 to 10 based on the BFP brightness/expression observed in their progeny (the brighter the higher the score). The best 6 founders for each transgene were isolated and outcrossed with tg(MN:GFP), with resulting clutches screened for embryos displaying blue fluorescence that were raised to adulthood to generate stable F1 responder lines.

Those F1 animals were further outcrossed with tg(UBI:iCRE) to evaluate the efficiency of this new conditional system. Interestingly, while all tg(UBI:loxBFP:mKate:INTRON_miRsmn1-4141) displayed bright red fluorescence (**table 1**), evidencing proper Lox-recombination of the tag-BFP cassette (**figure 4B**), none of the tg(UBI:loxBFP:dsRED:miRsmn1-4141) showed any detectable red fluorescence, suggesting proper recombination of the tagBFP cassette but no observable activation of the marker/RNAi block. Interestingly, although floxed tg(UBI:loxBFP:dsRED:miRsmn1-4141) did not display any red fluorescence, they did develop obvious traits of SMA, such as loss of motor function and premature death. Considering these preliminary observations, we hypothesised that the anti-*smn1* miRNAs were properly expressed but their processing/maturation hampered the expression of the red fluorescent marker, and decided to not follow up with those lines. On the contrary, all floxed tg(UBI:loxBFP:mKate:INTRON_miRsmn1-4141) displayed red fluorescence with brightness ranking similar to their unfloxed blue fluorescent counterparts, *i.e.* the brightest unfloxed blue fluorescent lines/embryos translated to the brightest floxed red fluorescent samples (**table 1**). We then selected the brightest F1 line, tg(UBI:loxBFP:mKate:INTRON_miRsmn1-4141)#1, also named tg(loxSMN) in this manuscript, to further analyse its ability to generate SMA-like larvae.

We previously showed that *smn1* loss-of-function (LOF) triggers motor neuron developmental defects, progressive loss of motor function and premature death as observed in patients^5^. We thereby tested if we were able to reproduce those phenotypes using this new system and generate a versatile model for research. To analyse those phenotypes, we performed a series of experiments in which we outcrossed tg(loxSMN) with tg(UBI:iCre) and sorted out embryos based on their fluorescence profile. Two controls were used, one with unfloxed embryos (BFP positive, mKATE negative), named “Unfloxed-CTR”, and another from a cross between tg(MN:GFP) and tg(UBI:iCre), named “iCre-CTR”. In term of motoneuron development, we found that floxed tg(loxSMN) developed significant abnormal ventral CaP projections with 2.55(±1.39) at 50 hours-post-fertilisation (hpf) when compared to controls iCre-CTR (0.55 ±0.75) and Unfloxed-CTR (0.6 ±0.75) (**Fig. 5 and 6A**). Demonstrating the specificity of this phenotype, injection of 250pg of human *SMN1* mRNA significantly reduced these defects with only 1.4(±1.35) abnormal CaP projection per side of animal. Floxed tg(loxSMN) also presented with a slight growth retardation with an average length (1379 ±45) significantly shorter than the controls iCre-CTR (1561 ±26) and Unfloxed-CTR (1580 ±21) (**Fig. 6A**). Injection of human *SMN1* mRNA also significantly rescued this growth retardation with an average of 1467(±20). Interestingly, these defects also translated into a progressive loss of motor function as presented below in **figure 7** as well as in premature death (**fig 6C**). Indeed, while 80% or more of both controls survived past the survey of 20 days-post-fertilisation (dpf), only 15% of the sorted floxed tg(loxSMN) animals survived more than 20 days-post-fertilisation (dpf); and none surviving until adulthood. Injection of 250pg of human *SMN1* mRNA did not rescue the premature death of the floxed tg(loxSMN), which was to be expected as injected mRNA usually gets degraded by, or before, 5 days-post-injection/fertilisation.

**Figure 5:**
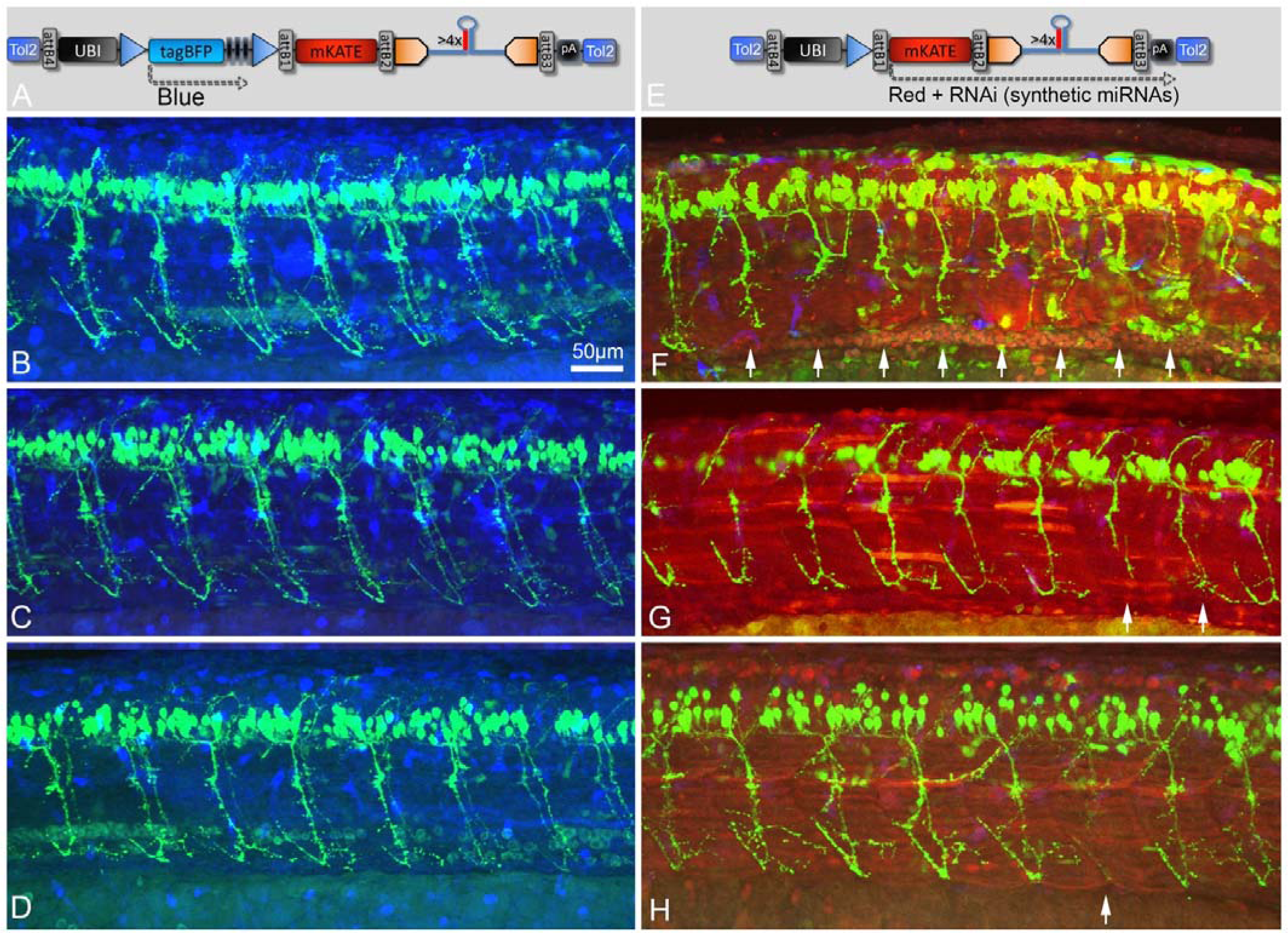
Schematics and representative images of 50hpf zebrafish larvae with unfloxed (A-B) or floxed (C-D) integrated RNAi transgene. A-B, In the absence of iCRE, the transgene remains unfloxed triggering BFP expression and not inducing any significant developmental defect in the analysed embryos (see figure 6 for quantification). C-D, In the presence of iCRE, BFP is excised and mKate expressed along with anti-*smn1* miRNAs, triggering SMA-hallmarks such as motoneuron defects. Images represent standard deviation projections from confocal z-stack acquisitions using three channels (merged) to detect eGFP (Green, motoneurons), tagBFP (Blue, ubiquitous expression) and mKate (Red, ubiquitous expression). White arrows indicate abnormal CaP motoneurons.

**Figure 6:**
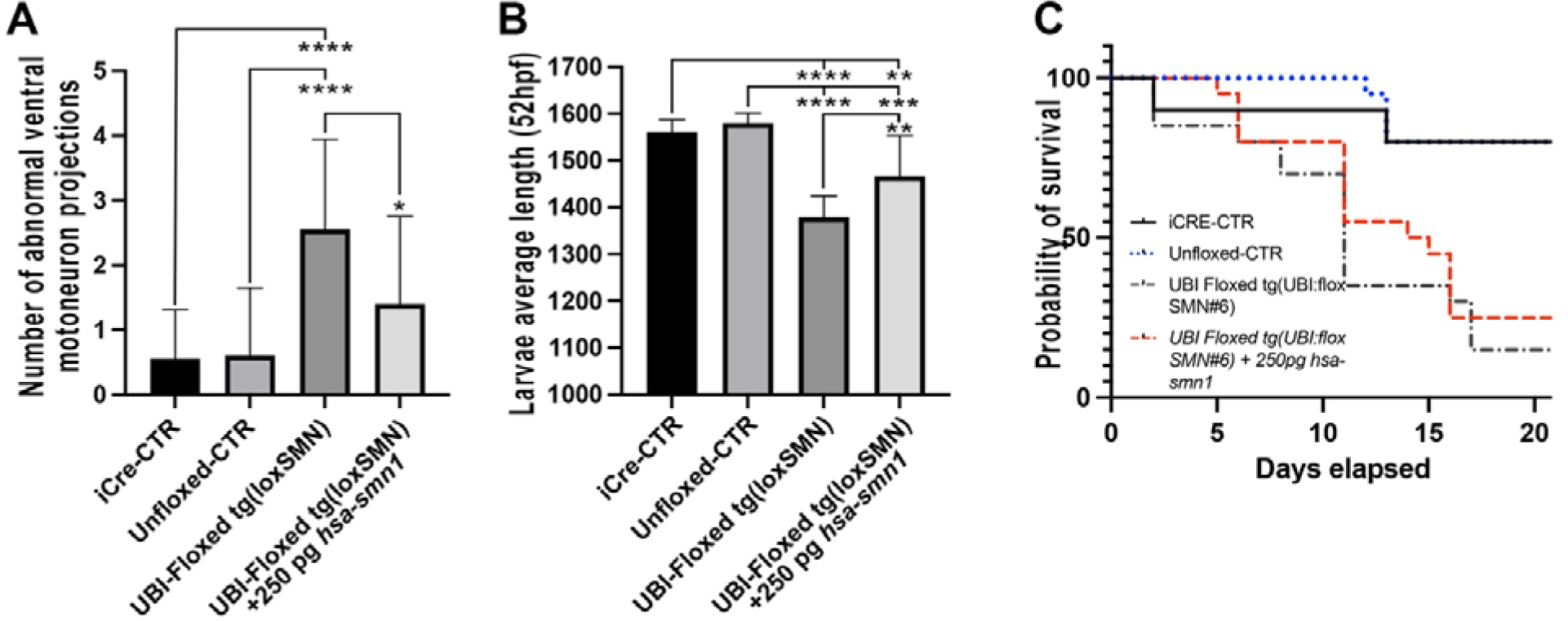
Phenotypic analysis of tg(floxSMN). **A,** Number of ventral motoneuron (CaP) abnormalities observed per side of 50hpf larvae. UBI-floxed tg(loxSMN) animals were also injected with *hsa-SMN1* mRNA to test the specificity of the observed phenotype. **B,** Animal size comparison between at 52 hpf (Arbitrary units). **C,** Survival assay (Kaplan Meier) for the different line and condition tested. Means of 20 larvae ±SD.

**Figure 7.**
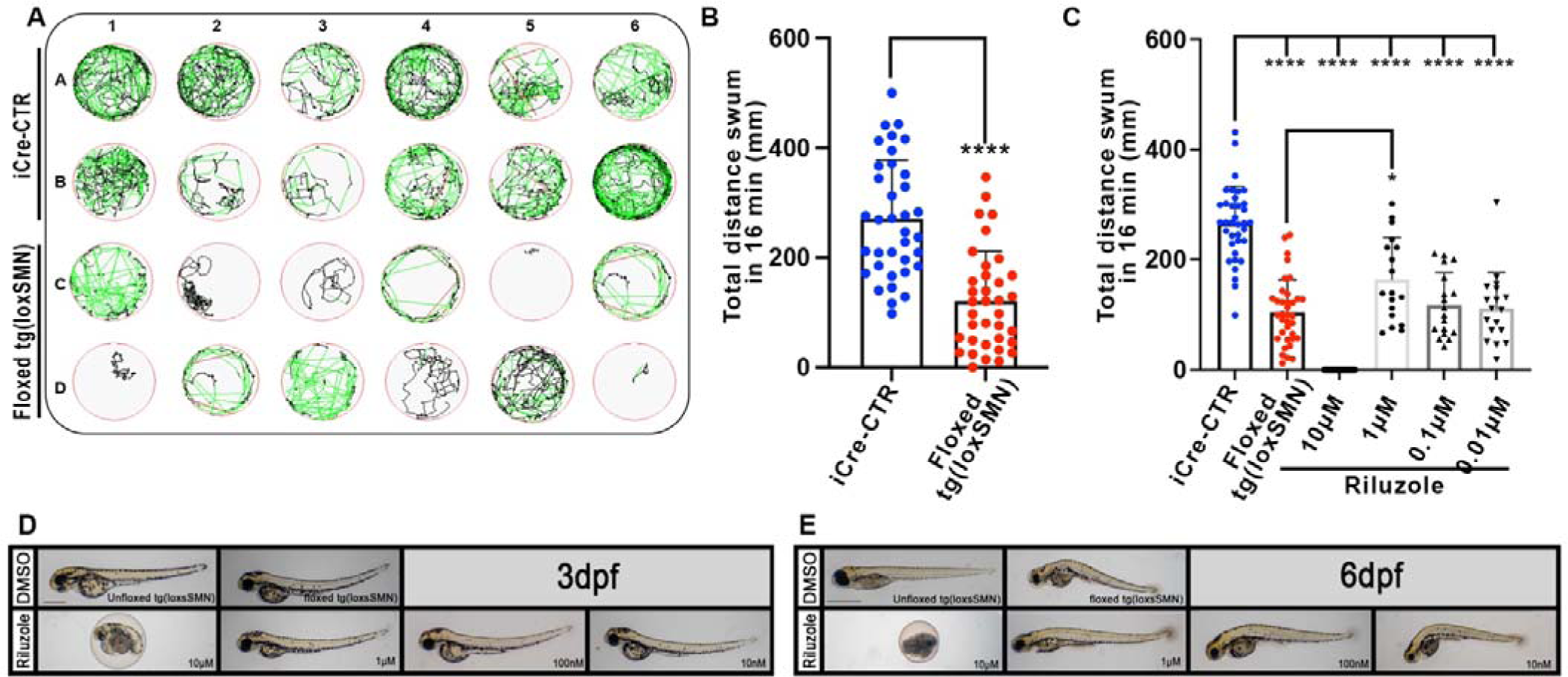
Conditional RNAi system suitable for large-scale experiments. Drug screening requires large number of “affected” samples that can be identified/genotyped prior to the screens. This could be a problem when one works with mutant lines. The presented iCre/Lox RNAi strategy offers the advantage of facilitating the generation of a large number of LOF-animals that can be identified prior to the screen thanks to their fluorescent profile (fig. S03). **A,** Swimming tracks of iCre-CTR and Floxed tg(loxSMN) after a 16min recording alternating 4min of light and dark phases. Black tracks represent slow speed (<2mm/sec), green tracks medium speed of (2-6mm/sec) and red tracks high speed (>6mm/sec). **B,** Average distance swum by 36 larvae over a period of 16min alternating 4 minutes of light and 4 minutes of dark. **C,** Effect of Riluzole on the swimming behaviour of floxed tg(loxSMN) animal. Results are represented as means ±SD with dots representing a single data point or well.

All in all, these results demonstrated that our Cre/Lox miR-delivery/RNAi technology is effective and that we successfully generated a conditional genetic system able to produce healthy RNAi-carriers that can be “ranked” and easily maintained thanks to the fluorescence associated with the unfloxed state. We showed that, while asymptomatic/silent, these lines can be easily activated to recapitulate strong human disease hallmarks, such as the SMA defects and premature death presented here. Interestingly, these silent RNAi-responder transgenics can be activated not only by transgenic cross with Cre-driver lines but also by injection of either Cre/iCre mRNA or Cre protein^16^. In a side study, we have already worked at defining the recombinant effect of different Cre mRNAs and protein versions as well as dosage effects for controlling this new genetic miR-delivery system^16^.

### Versatile genetic system for drug discovery or large-scale experiments

Drug discovery using the zebrafish animal model is becoming popular. However, it is often difficult to develop models of human diseases that would be compatible with large-scale (high throughput/content) screening. <u>First</u>, one need to establish a model that develops robust and early phenotype, *i.e.* ideally taking place in the first 2-7dpf. Interestingly, here we generated a model of choice. The presented floxed tg(loxSMN) develops progressive loss of motor function (**fig 7A and 7B**) and premature death (**Fig. 6C**) that can be used as readouts in a screen. Animals start to demonstrate a decrease in their spontaneous swimming behaviour starting from 5dpf and with a clear abnormal pattern at 6dpf, as demonstrated in **fig. 7A** and **7B**. Our assay consisted of an automatic record of the larvae swimming tracks for 16 minutes with alternating light and dark phases. In these conditions, while control iCre-CTR swam for an average of 271mm (±107mm), floxed tg(loxSMN) swam significantly less with an average of 121mm (±91mm). Although relatively heterogeneous, this readout could be used to establish a screen, looking for compounds improving/rescuing those motor defects. <u>Second</u>, and most importantly, to be able to run a drug screen or a large-scale experiment, one should be able to produce a large number of affected animals that can be identified before the assays or readouts. For example, when one uses mutants with early phenotype, usually only heterozygotes could be maintained for breeding with only 25% of the resulting embryos producing early phenotype (**fig. S03A**). One has to either be able to genotype the embryos before the experiments, or deal with 75% data points that would bias the readouts/results. In contrast, here, one can sort the embryos based on their fluorescent expression prior to the screen (**fig. S03B and S03C**). Moreover, we found that crossing female tg(UBI:iCRE) drivers with male RNAi responders, led to 100% conversion of the RNAi lox cassette, whether the female tg(UBI:iCRE) driver was heterozygous or homozygous, strongly facilitating the set-up of screening plates. We hypothesised that this was due to maternal deposition of iCre-mRNA and iCRE-protein into the embryos^16^. Demonstrating the value of this genetic system and approach for drug screening, we tested serial dilutions of the neuroprotective drug Riluzole in 24-well plates, from 100µM down to 100nM. We found that 100µM was lethal while 10µM slightly improved motor functions of floxed tg(loxSMN) animals at 6dpf (**fig. 7C**). No further effect was observed on either motoneuron development or premature death, suggesting unfortunately a poor therapeutic outcome on the presented zebrafish SMA model.

### miR-delivery genetic system enabling tissue-specific studies

We further tested the ability of the presented system to conduct tissue-specific studies. We generated transgenic lines using tol2 iCre-transgenes designed to drive pan-neuronal- (HuC promoter), motoneuron- (HB9 promoter) or striated muscle- (503UNC promoter) expression. Each transgene co-expresses an eGFP marker under the control of a crystallin promoter, enabling straightforward tracking of the carriers and downstream lines. Unfortunately, although we isolated stable transgenic lines for each transgene, none of our tg(HB9:iCre) lines successfully triggered observable lox recombination. For that reason, we could not generate motoneuron-specific data in this manuscript. However, we successfully generated a functional tg(HuC:iCRE) line and showed that pan-neuronal Lox-recombination in tg(loxSMN) triggered motoneuron defects similar to the one observed in ubiquitous lines (**fig. 8**). Surprisingly, these defects were not as pronounced as in the ubiquitous condition, with only 1.3(±1.4) abnormal CaP branch at 50hpf, significantly different from 2.45(±1.4) at 50hpf in the ubiquitously floxed lines (**fig. 8B**). This difference may be explained, at least in part, by the fact that the gene *smn1* is ubiquitously expressed and that a certain amount of SMN protein should already be present in the early differentiated neurons before activation of the miR-mediated *smn1*-knockdown following iCre transcriptional activity. This feature could certainly represent a limitation of this technology when one desires to work tissue-specifically. In contrast, although striated muscle expression of the anti-*smn1* miRNAs triggered significant early motoneuron defects (0.9±0.8 at 30hpf), there was no further evidence of significant permanent motor neuron phenotype at later stage (0.65±0.9 at 50hpf). All conditions did significantly impact the survival of the animals. However, it is noticeable that most 503UNC-animal deaths happened at around 11dpf and seem to be related to an absence of an inflated swim bladder. All in all, those data demonstrate the potential of this system to drive inducible and/or tissue-specific gene-silencing experiments. This controllable genetic system should also be very useful for microRNA basic research studies by allowing versatile control and expression of endogenous miRNA(s) of choices.

**Figure 8.**
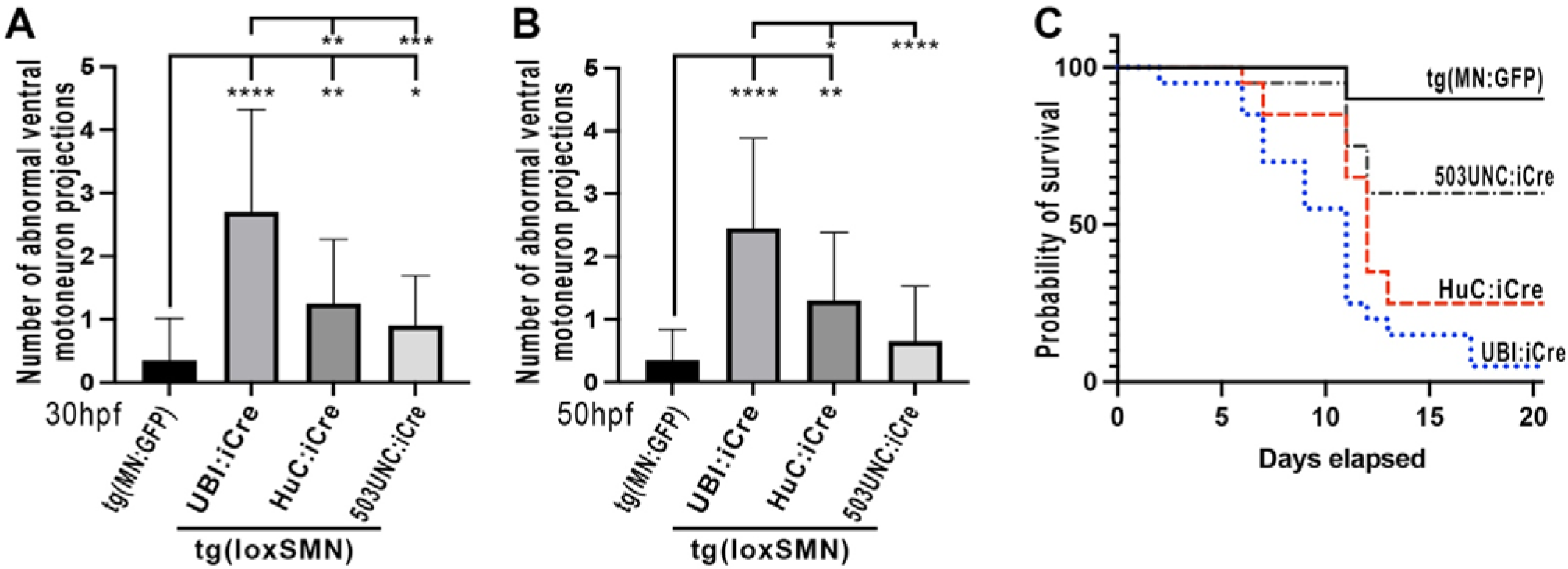
The RNAi Cre/Lox system allows tissue-specific gene(s) silencing. Numbers of ventral motoneuron (CaP) abnormalities observed per side of 30hpf **(A)** and 50hpf larvae **(B)**. **B,** Survival assay (Kaplan Meier) for the different lines tested. UBI. Ubiquitous promoter. HuC, Pan Neuronal promoter. 503UNC, Striated muscle promoter. Means of 20 larvae ±SD.

### Rapid cell-specific miR-delivery/RNAi studies

As presented in Figures 2 and 3 above, the ability to track the injected miR-delivery/RNAi transgenes opens the door to rapid mosaic cell-specific experiments. Following injection, each cell presenting transcriptional activity of the injected transgene will present fluorescence and could be compared to their non-expressing counterparts within the same animal. To optimise the injection mix for such an approach, we constructed a motoneuron-specific anti-*smn1* construct combining 106_p5E-HB9, 010_pME-mKate2, 61_p3E_INTRON_*smn1-4141* with a 394_pDestTol2pA2 destination clone. The final construct was named 83_HB9_mKate_smn4141. We injected this construct in one-cell stage tg(MN:GFP) animals at different concentrations, ranging from 25pg to 200pg final, complexed or not with transposase (**fig. 9**). Our experiments suggest that a concentration of up to 50pg of plasmid per cell is well tolerated while toxicity is observed starting from 100pg onward (**fig. 9B**). We then quantified the number of animals presenting at least 1x mKate-positive CaP motoneuron (evidencing the activity of the miR-delivery transgene, **fig 9A arrowheads**) and plotted the different percentage obtained in **figure 9C**. Interestingly, our data suggest that the addition of transposase (promoting transgene integration) did not seem to greatly impact the percentage of positive animals per injected batch (**fig 9C**) or the average number of positive expressing cells per injected animal (**fig 9D**). While 25pg of plasmid alone led to a batch with 27.4% animals presenting at least 1x mKate-positive CaP -with 2.91+/-2.73 positive CaP-, the addition of transposase mRNA (25pg) did not dramatically change this ratio with only 34% of positive animals -with 3.92+/-3.3 positive CaP-. However, the injection of 50pg of plasmid alone led to 72.6% of positive animals presenting an average of 4.3+/-3.2 positive CaP per positive animal. From 50pg, the addition of transposase or a higher plasmid dose led to significant malformations that would interfere with the experiments (**Fig. 9B**). Interestingly, the expression of the transgene (with or without transposase) was detectable long after injections with constant pattern at least up to 6dpf. Those results suggest that a dose of 50pg without transposase would be a range of choice for conducting mosaic cell-specific experiments.

**Figure 9.**
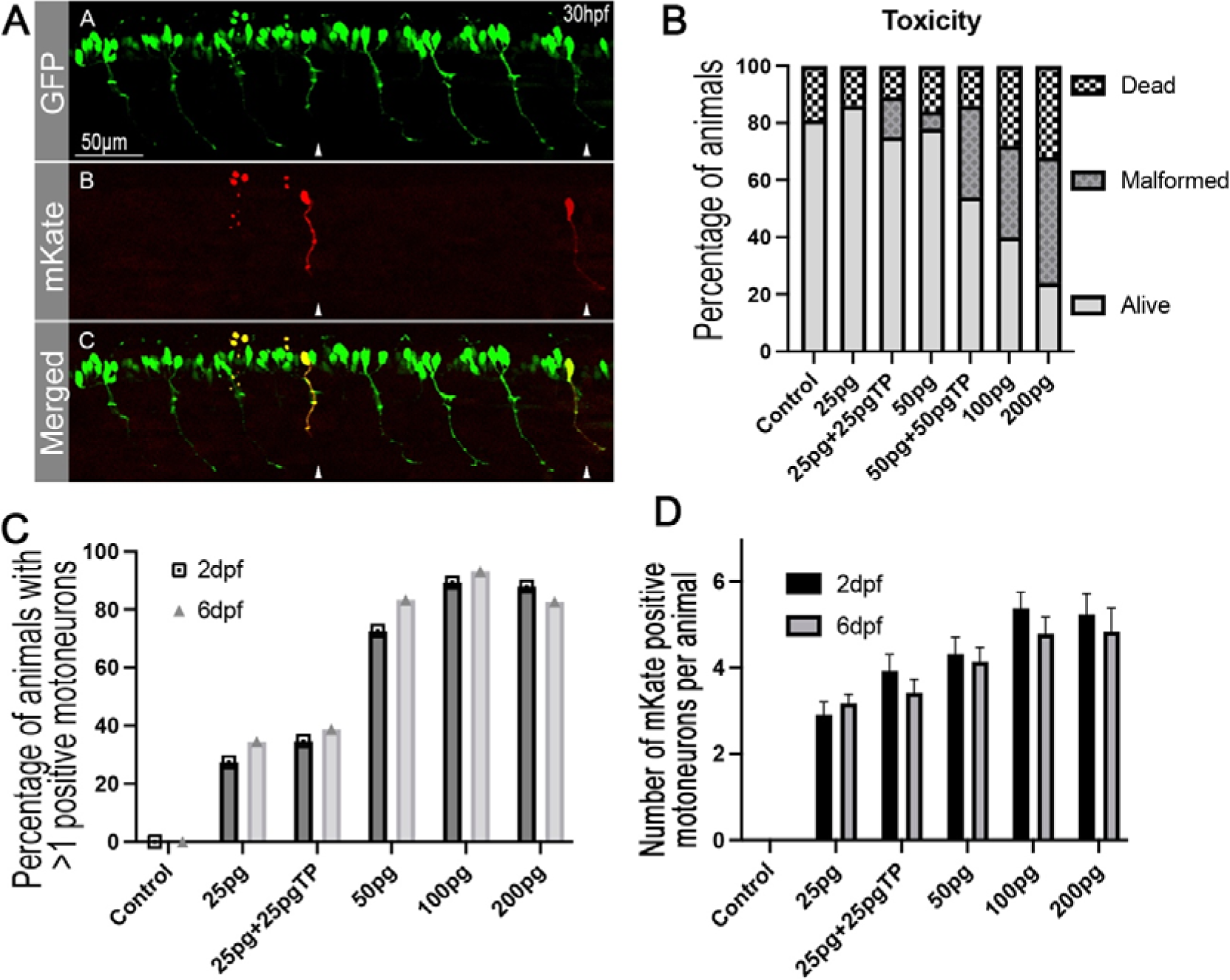
Cell-specific miR-delivery/RNAi in zebrafish does not require transposase. We tested the ability of the presented miR-delivery system to enable rapid cell-specific experiments. The approach is similar to the method conducted in Figures 2 & 3 but aimed at evaluating the effect of different injection mix. **A,** Schematic example of the expression obtained following the injection of a motoneuron-specific miR-delivery construct (83_HB9_mKate_smn4141) within the cell of one-cell stage tg(MN:GFP) animals. Arrowhead indicating mKate positive motoneurons with positive transgene expression. **B,** Titration/Toxicity test of tol2-construct injections (83_HB9_mKate_smn4141) for triggering cell-specific expression. **C,** Number (Percentage) of animals presenting at least 1x mKate positive motoneuron. **D,** Average number of mKate-positive CaP motoneurons per positive animal. 80 embryos per condition have been analysed. These results suggest that a dose of 50pg without transposase may be ideal for setting up cell-specific approaches and studies.

## DISCUSSION/Conclusion

We have previously demonstrated that miR-mediated gene knockdown could be a powerful tool for zebrafish genetics and that this technology has the potential to be a versatile complement to CRISPR/Cas9 approaches^21^. Those approaches could also ease the studies of endogenous microRNA-biology by offering a versatile genetic system for transgenic delivery. Unfortunately, those tools were suffering from important limitations that were hampering their potential. <u>First</u>, let alone multi-genic silencing, to achieve proper single-gene knockdown, multiple synthetic miRs must be expressed simultaneously which compromises the co-expression of a fluorescent marker/tracker, strongly reducing the interest in this approach. <u>Second</u>, when knockdown of the gene(s) of interest triggers significant phenotype/defects, it is often hard to generate and maintain stable transgenic carriers with strong expression.

Here, we optimised this gene-silencing and miR-delivery approach and established a new and mature genetic system for zebrafish. Based on our analyses, this genetic system does no longer suffer from limitations due to the processing of the synthetic miRs, and one can now chain multiple miR-delivery/RNAi cassettes without taking the risk of losing the coupled fluorescent tracker/marker. In theory, any combination of microRNAs can now be simultaneously expressed, and quantitatively monitored via the associated fluorescence. Any gene of interest can now be targeted, and multi-gene knockdowns should also be easily achievable. Although this approach may not be able to surpass the knockout efficiency and reliability of the recent CRISPR/Cas9 methodologies, the ability to track cells, tissues and organisms prior and during the analysis is of unique value and flexibility for research. In addition, we are delivering here a conditional genetic system that proves to be powerful enabling the generation of healthy carriers that can now be easily maintained. Importantly, even when not activated, this system offers the advantage of enabling the selection and monitoring of transgenic lines/carriers with high transcriptional activity of the “silent” miR-delivery/RNAi transgene, thanks to the BFP/blue expression (different sets of fluorescence markers could be used). Another important advantage is the ability to drive the activity of the miR-delivery/RNAi cassette spatiotemporally, opening the door to cell- and tissue-specific investigations, an approach that is still unmatched today with the zebrafish model. Moreover, the potentiality to either use transgenic Cre-driver or injectables (Cre mRNA or protein)^16^ offers great versatility for the investigators, allowing for instance to mix different responder-RNAi without performing complicated and lengthy crosses. We also demonstrated that, using this method, one could easily produce a large number of embryos that would not require a pre-screen before the experiments (**fig. 04 and S04**), offering greater flexibility for research and large-scale experiments. Last but not least, this method and material should be of high value to express and study endogenous microRNAs biology. For example, here to validate our tools, we did over-express an endogenous miRNA, *i.e. miR137*. Although we expressed it in a tissue where it is not present naturally, this approach demonstrates that, using this technology, one can express any miRNA of choice to study its role in a whole organism context.

## CONFLICT OF INTEREST

None to declare.

## AKNOWLEDGMENTS

This work was supported by grants from the Australian National Health and Medical Research Council (NHMRC) Project Grant No 1165850 to JG, The Rebecca Cooper Medical Research Project Grant No PG2019405 to JG and a CureSMA of USA grant to JG. JG was also supported by an NHMRC Emerging Leader Fellowship No 1174145.

## METHODS

### Zebrafish Maintenance

Adult zebrafish and embryos were maintained by standard protocols approved by the University of Queensland and Griffith University Animal Ethics Committee. Ethic approval AE213_18/AE213_18 and GRIDD/11/22/AEC. Wild-type lines used in these studies are TAB background.

### Cloning of p3E-RNAi

We first cloned a p3E clone incorporating the rabbit beta-globin intron presented here (“Kaloop” plasmid)^12^. The intronic sequence along with a SV40 signal was amplified using forward, Rabbit_GI_F1_p3E (ggggacagctttcttgtacaaagtggCTCGACCGATCCTGAGAACTT), and reverse primer, SV40_late_pA_R1_p3E (ggggacaactttgtataataaagttgCCACACCTCCCCCTGAAC). The generated PCR product was further recombined into pDONRp2R-p3 using Gateway LR clones following manufacturer instructions. Final plasmid was named 303_p3E_GI_PA in our database. We then introduced into this 303_p3E_GI_PA our previous RNAi cassette from plasmid pME-RNAi641^5^ using an “In-fusion (Takara)” cloning strategy. We digested 303_p3E_GI_PA using DraI (middle portion of of the beta globin intronic sequence) and purified it. In synergy, we amplified the previously published RNAi cassette using In-fusion compatible forward (TTGTAACGAATTTTTcgtcgatcgtttaaagggagg) and reverse-primer (TTGTAACGAATTTTTcgtcgatcgtttaaagggagg). We conducted an In-fusion cloning reaction (In Fusion HD kit, Takara) following the manufacturer instructions to introduce the RNAi cassette into the digested intronic sequence. The kanamycin gene resistance sequence in the resulting clone presented an XhoI site that would interfere with the chaining ability of the RNAi cassette. We then removed it using a site directed mutagenesis using Phusion High Fidelity DNA Polymerase and the following forward and reverse primers (CGATCGCGTATTTCGcCTCGCTCAGGCGCAA and TTGCGCCTGAGCGAGgCGAAATACGCGATCG). Reaction conducted was as follow, first denaturation step at 98°C (15sec) followed by 18 cycles of 98°C(15sec)/65°C(30sec)/72°C(1min) and terminated by an elongation step 72°C(5min). A sample of 5µl was run on a gel to verify the presence of a single appropriately sized band. The remaining reaction mix was then treated with 0.3µl of DpnI for 10 min at 37°C. A volume of 2µl was finally used to transformed competent cells and plated on plates supplement with kanamycin. The resulting colonies were screened by miniprep and sequencing. Final plasmid was named p3E-IntronRNAi (addgene #163380) also recorded as 61-p3E-IntronRNAi in our database.

### Artificial/synthetic *pri-miRNA* design, assembly, and insertion into p3E-IntronRNAi

In order to design a *pre-miR* to insert into the p3E-IntronMIR, the minimal 3’UTR region of the gene(s) of interest should first be identified and confirmed. Targetscan Fish (http://www.targetscan.org/) provide useful resources in addition to traditional online databases such as *Ensembl*. However, we strongly recommend sequencing the 3’UTR of interest to confirm the integrity of the identified sequence on the mRNA(s) to deal with any potential polymorphisms. Once confirmed, target locations should be identified manually or using online tools. We recommend the miRNA-design tool provided by Invitrogen (www.genscript.com/design_center.html), which generates oligos compatible with p3E-RNAi, thereby ready for order. For a manual design, follow the guidelines in **figure S05**. Other possible online tools are described in the original method here^5^. After the design, order the top and bottom *pre-miRNA* oligos using your favourite oligos-provider, and resuspend them at 200mM for storage. For generating the *pre-miR* to insert into BsmBI-digested p3E-IntronRNAi, anneal 5µl of these 200mM Top and Bottom RNAi oligonucleotides in a 20µl reaction including 2µl of NEB2 10x buffer. The mixture is heated at 95°C for 5min in a thermocycler and left in the machine to cool down for another 30min. Samples are then vortexed and briefly spun down before being diluted 5,000-fold in water at room temperature. Diluted final *pre-miRNA* samples are stocked at room temperature until ligation into BsmBI-digested p3E-IntronRNAi. Discarded after use. For inserting those *pre-miRNA*, p3E-IntronRNAi is first digested by BsmBI and gel extracted. Resulting linearised plasmid and pre-RNA are ligated as follows: Mix 10ng of linearized p3E-IntronRNAi with 4µl of the 5,000-diluted *pre-miR* and ligated for 1hour at room temperature before transforming Max efficiency bacteria and plate on kanamycin. The next day screen your colonies with forward (46_F_RNAi aagggaggtagtgagtcgac) and reverse (47_F_RNAi ctagatatctcgagtgcggc) primers. We also recommend using those primers for sequencing. Note that BsmBI-digested p3E-IntronRNAi can be stored for several years after gel extraction and purification. Following the insertion of the annealed *pre-miR*, you will have generated functional *pri-miR* that can be chained as presented in **figure S02** or as described below.

### Chaining of RNAi cassettes

Several *pre-miRNA* have been used here, such as *pre-miR-137* and *pre-miRsmn1-4141*, have previously been cloned and described^5,14^. For the 1x *miR137* constructs, previous *pri-miR-137*^14^ have been gel extracted from UAS:YFPs:137 using BamHI/XhoI digestion, and inserted into p3E-IntronRNAi (also digested with BamHI/XhoI, gel extracted and purified), with the final plasmid named “p3E-Intron_miR137_1x”. *p3E-Intron_miR137-4x* (4x *pri-miR-137* repeats) was further generated through chaining of 4x *pri-miR-137* cassettes as presented in **figure S02A**. For generating *p3E-Intron-smn1-4141* used in this study, we gel-extracted the *pre-miRsmn1-4141* cassette from 641-dsRED-smn1-4141, published here^6^, using BamHI/XhoI digestion. We further inserted this *anti-smn1* cassette into p3E-IntronRNAi for downstream cloning presented below. This p3E was named p3E-Intron-smn1-4141.

### Transient experiments via RNAi-transgene injections

To validate the intron efficiency, we first generate Tol2-transgenes using a LR-reaction combining 394-pDestTol2pA2, 104_p5E-503UNC^6^, 383-pME-EGFP (<u>tol2kit</u>) and the corresponding p3E with or without intron. We then injected 50pg of each plasmid, named 153_503UNC-eGFP_1xmiR137, 154_503UNC-eGFP_4xmiR137, 155_503UNC-eGFP_INT_1xmiR137, 156_503UNC-eGFP_INT_4xmiR137 and controls along with tol2 transposase mRNA into 1-cell staged embryos (**figure S03**). These embryos were from an outcross of Tg(UBI:mCherry:SP137) fish ubiquitously expressing mCherry tagged with a custom 3’UTR recognised but *miR-137*^14^. We imaged the animals (muscle fibres) at 4dpf using confocal microscopy to ensure proper quantification of knockdown. We processed and analysed the recordings using Fiji. To quantify pixel intensity, we used “plot profile” to extract gray values across a line drawn across regions of interest (ROI). We plotted red fluorescence as a percentage and green fluorescence as raw values and generated linear regression curves using PRISM9.

### Generation of Driver lines

Plasmid UBI:iCre, HuC:iCre, HB9:iCre and 503UNC:iCre have been produced using a 2-way Gateway LR reaction using manufacturer instructions. For all final plasmids, a custom R4-R2_destination clone named 1456-pDEST-minTol2_R4-R2_Cryst-eGFP (Addgene #171795, carrying a selection eGFP marker under the control of the Crystallin promoter) was mixed with our 469-pME-iCre (Addgene #171792) and either with 101_p5E-Ubiquitin, 102_p5E-HuC, 106_p5E-HB9(3.3kb) or 104_P5E-503UNC. Each plasmid was injected into TAB fish at 25ng.µl complexed with 25ng.µl of transposase-mRNA for promoting DNA integration. Around 200 animals were injected and around 60 larvae were selected at 3dpf based on visible green fluorescence expression in their eyes (Lens). Larvae were raised until adulthood and screened for F0 founder/carrier. Three F0 adults were selected based on strong eGFP Lens-expression in their F1 progeny. The F1s were screened and raised until adulthood to be outcrossed with responder lines to test their ability to trigger Lox-recombination. Only one line was further maintained based on the validation/efficiency of the associated Cre activity. It is noteworthy that we failed to identify an efficient/strong HB9-driver.

### Cloning of p5E-UBI:LoxBFP

We first amplified tagBFP from our 15_pME_tagBFP stock plasmid using primers 101-BamHI-Flox-BFP-ForwRight (acagggatccaagcttataacttcgtatagcatacattatacgaagttatccggtcgccaccatgagcgagctgattaagg) and 102-BFP-NotI-Rev (AATTggatccgcggccgctttaattaagcttgtgccccagt). The primers were designed to insert a Flox sequence in 5’ of the tagBFP sequence, the overall amplicon being flanked by BamHI and NotI sites. The PCR product was further digested using BamHI along with tol2kit 101_p5E-Ubiquitin plasmid (Addgene #27320). Both were purified and 101_p5E-Ubiquitin was dephosphorylated using NEB Antarctic phosphatase following manufacturer instructions. T4 ligation was conducted overnight at 12C and transformed using MAX Efficiency DH10B Competent Cells (Invitrogen #18297010). Bacteria were grown on kanamycin resistance plates overnight at 37°C. Ten colonies were amplified in 4ml of LB supplemented with kanamycin. Plasmids were isolated through miniprep (PureLink Quick Plasmid Miniprep Kit) and sent for sequencing to confirm orientation of the insert and integration of the Lox_tagBFP cassette. The intermediate plasmid was named p5E-UBI_Lflox_BFP. We further digested it using NotI, purified and dephosphorylated it. In parallel, we gel extracted the 3x polyA sequence present on plasmid pENTR5’_ubi:loxP-EGFP-loxP (Addgene # 27322) and inserted it in the NotI-dephosphorylated p5E-UBI_Lflox_BFP plasmid through T4 ligation. 10 colonies were selected for miniprep as presented above and sequenced for selecting a clone with proper orientation. Final plasmid was named p5E-UBI-loxTagBFP (Addgene #172518) and recorded in our database as 50_p5E-UBI-loxTagBFP.

### Cloning of UBI:loxBFP:dsRED:miRsmn1-4141 andcUBI:loxBFP:mKate:INTRON_miRsmn1-4141

UBI:loxBFP:mKate:INTRON_miRsmn1-4141 was cloned using a Gateway LR-reaction mixing destination p5E-UBI-loxTagBFP (Addgene #172518) with our 010-pME-mKate2, p3E-Intron-smn1-4141 and destination clone 397-pDESTtol2pA_Cryst-cherry. Reaction was transformed using Max efficiency bacteria and selected on ampicillin plate. A PCR screening was performed, and two positive clones were sent for sequencing. UBI:loxBFP:dsRED:miRsmn1-4141 was generated through a similar cloning approach but mixing p5E-UBI-loxTagBFP with 641-dsRED-smn1-4141 and destination clone 1457-pDEST-minTol2_R4-R2_Cryst-cherry.

### Generation of Responder lines

Tol2-destination clones UBI:loxBFP:dsRED:miRsmn1-4141 and UBI:loxBFP:mKate:INTRON_miRsmn1-4141 were injected into one-cell stage tg(MN:GFP) embryos, with 30ng.µl of corresponding DNA complexed with 25ng.µl of transposase^5,18^. Injected animals were screened at 3dpf based on blue mosaic expression and/or red fluorescence expression in their eyes. Embryos were raised until adulthood and outcrossed with wt to identify F0 founders which integrated the respective conditional RNAi transgenes, which was evidenced by both ubiquitous blue fluorescence expression and eye-specific red fluorescence emission. We selected the best 6 lines that we ranked based on their apparent blue fluorescence brightness (**table 01)** and generated respective F1 stable lines. Once sexuality matured, those F1 lines were further outcrossed with tg(UBI:iCre) to characterise their ability to Flox/recombine and to express the conditional miR-cassettes, which was evaluated by observing and ranking red fluorescence emission (**table 01)**.

### Microscopy and motor neuron observation

To quantify motor neuron abnormalities and other phenotypes below, we generated synchronised embryos for the different conditions tested. Corresponding adult male and female animals were separated the day before mating to be merged in fresh water the next day. Animals were monitored and left mating for 1h. The resulting embryos were considered “synchronized” and collected in E3 medium without methylene blue. When appropriate, the embryos were injected with *hsa-SMN1* mRNA (injection in yolk of one-cell stage). The day of analysis, animals were dechorionated if required, anaesthetized in tricaine and observed sideways under a fluorescent microscope, unless stated otherwise. Only CaP-axon projection abnormalities were scored. One point was attributed to every motor axon that showed defects as abnormal branching, abnormal length or absence of projection. Scores were compared in PRISM 9 using t-tests.

### Survival Assay

Synchronised embryos were collected in E3 medium. When appropriate, the embryos were injected with human *SMN1* mRNA. Animals were sorted out at 1dpf based on their fluorescence profile and cultured in petri dishes for monitoring until 10dpf. Paramecia were added to the medium starting from 6dpf and animals were transferred into beakers starting from 10dpf. Embryos/larvae were then counted each day to record death ratios over a period of 20 days.

### *hsa-SMN* mRNA production

*hsa-SMN* mRNA was produced as previously described^5^. Briefly, pCS2+:hsaSMN1 was linearized with NotI, purified and used with a mMESSAGE mMACHINE SP6 transcription kit (Ambion), following the manufacturer’s protocol. *hsa-SMN* mRNA was purified using a MEGA clear kit (Ambion) following the manufacturer’s protocol, aliquoted and stored at −80°C. *hsa-SMN* RNA (250 pg) was injected into the yolk of one-cell-stage embryos when required.

### Behavioural experiments and Drug screening

Behavioural analysis was performed using the Zebrabox (Viewpoint) following the manufacturer’s instructions. Zebrafish larvae were distributed in 24-well plates filled with 500µl of E3 medium (one larva per well, +/- Riluzole as described or DMSO 1%). Following the plates setup, all samples were placed at 28°C for a minimum of one hour prior to run the analysis. The assays consisted of automatic recording of the larval swimming tracks/behaviour during 16mins and under alternating phases of 4 minutes light and 4 minutes darkness. At the end of the experiment, each larva was monitored to exclude from the data potential dead animals. Data were exported and processed using Excel/Prism.

### Cell-specific experiments

To test our material for cell-specific experiments, we generate an anit-*smn1* motor neuron specific construct. We combined 106_p5E-HB9, 010_pME-mKate2, 61_p3E_INTRON_*smn1-4141* with 394_pDestTol2pA2 to generate the final injectable construct 83_HB9_mKate_smn4141. We then injected different concentrations as described (+/- transposase) within the cell of one-cell stage tg(MN:GFP) embryos (**figure 09**). We analysed the animals at 1dpf, 2dpf and 6dpf to assess toxicity, fluorescent expression and motor neuron development patterns. Data were exported and processed using Excel/Prism.

## Notes

### Competing Interest Statement

The authors have declared no competing interest.

