## Supplementart Files for "Cre-Lox miRNA-delivery technology optimized for inducible microRNA and gene-silencing studies in zebrafish"

### Supplemental material

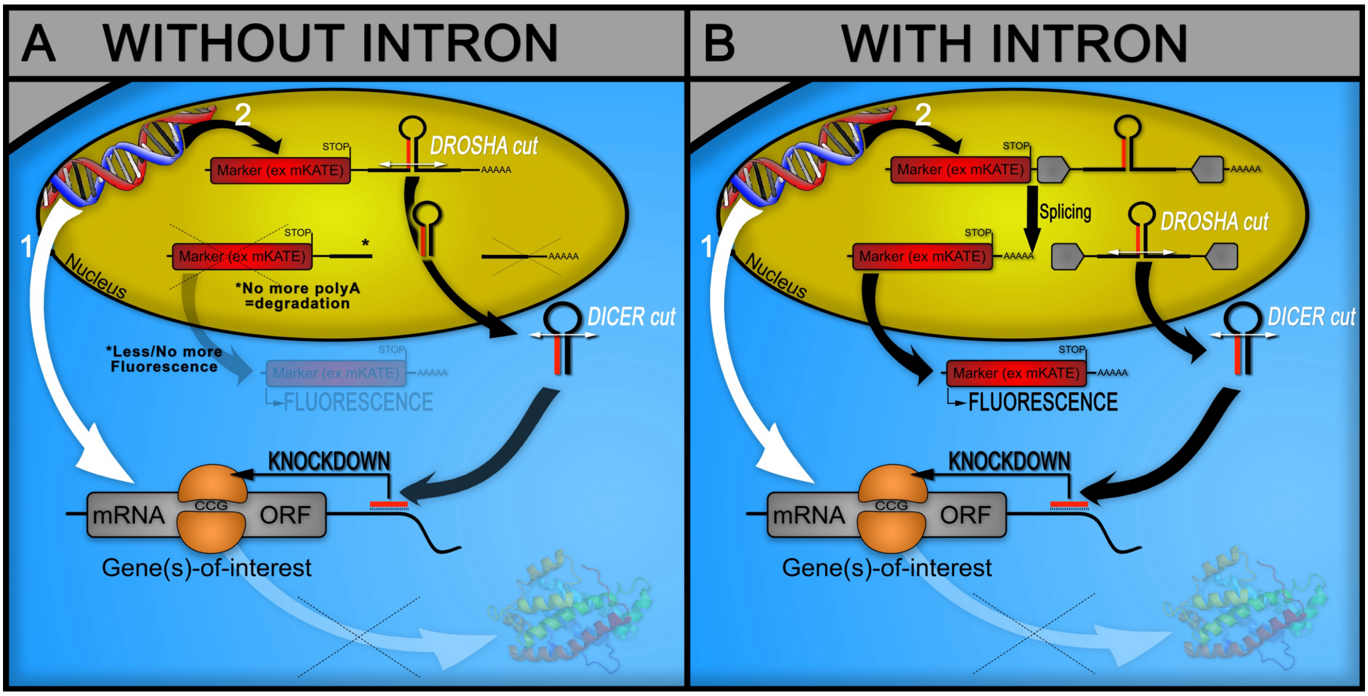


**Figure S01. Schematic representation of transgenic miRNA-delivery technique in zebrafish and the benefit of using intronic sequence for proper processing. A,** Without Intron, the miRNA-delivery cassette will be included in a mRNA along with a fluorescent marker. Unfortunately, the maturation of the miRNA will involve cuts that will lead to the denaturation of the mRNA due to the loss of its polyA tail. Consequently, the translation of the fluorescent marker and the maturation of the synthetic miRNA will work competitively, with the brightness of the marker not being proportional/correlated to the expression of the *miRs*. The fluorescence would be completely lost when using 3x or more *pri-miR* repeats. **B,** The introduction of an intronic sequence solves this limitation by inducing splicing of the *pri-miR* cassettes prior to Drosha cuts, leaving the host mRNA intact/stable and enabling co-expression of the attached fluorescent marker.

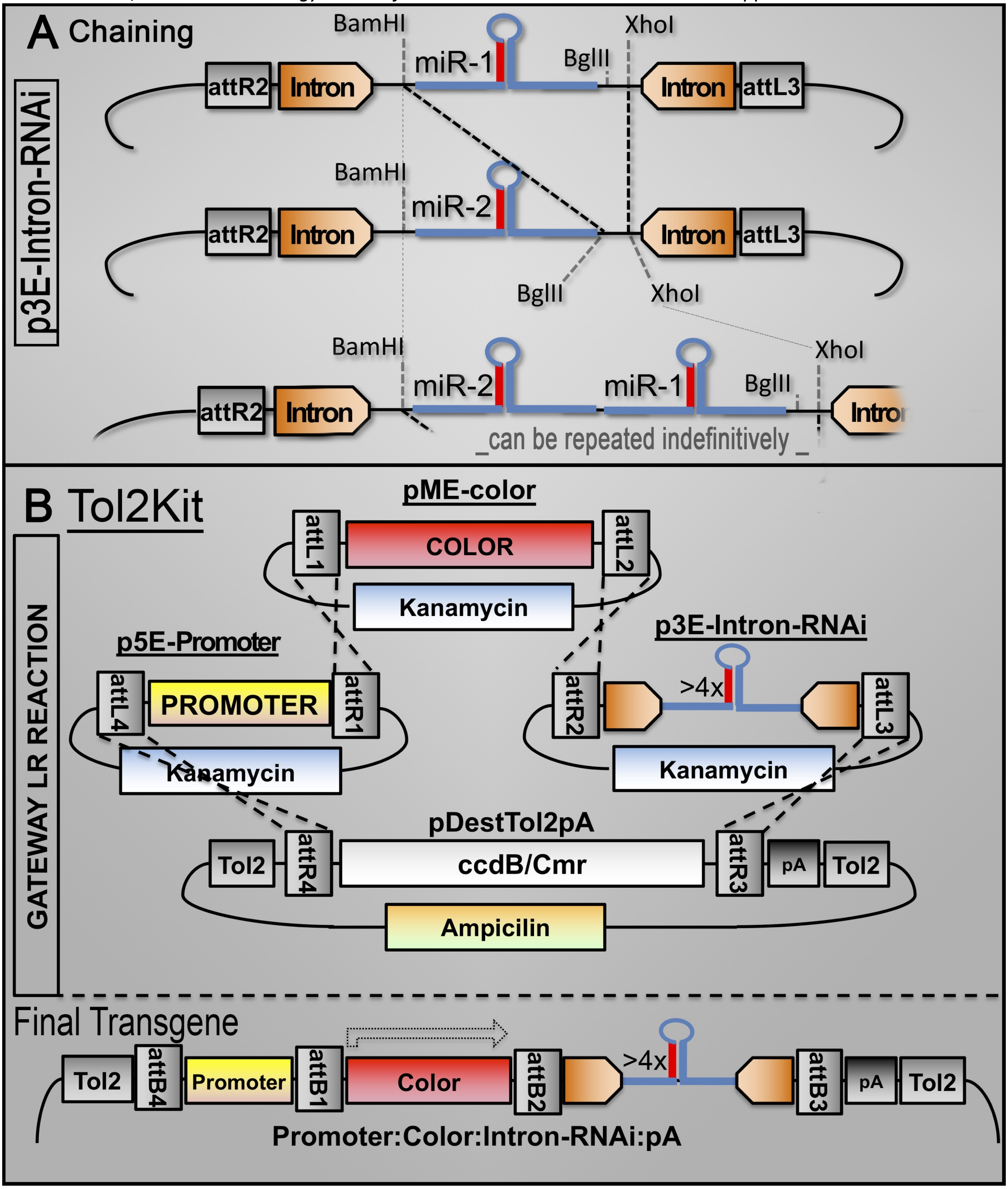


**Figure S02. Schematic representation of p3E-Intron_miR_RNAi cloning procedures. A,** The P3E plasmid is optimised for easily chained multiple *pri-miR* cassettes as concatemer using digestion reactions that can be repeated through unlimited rounds. **B,** Typical Tol2kit reaction.

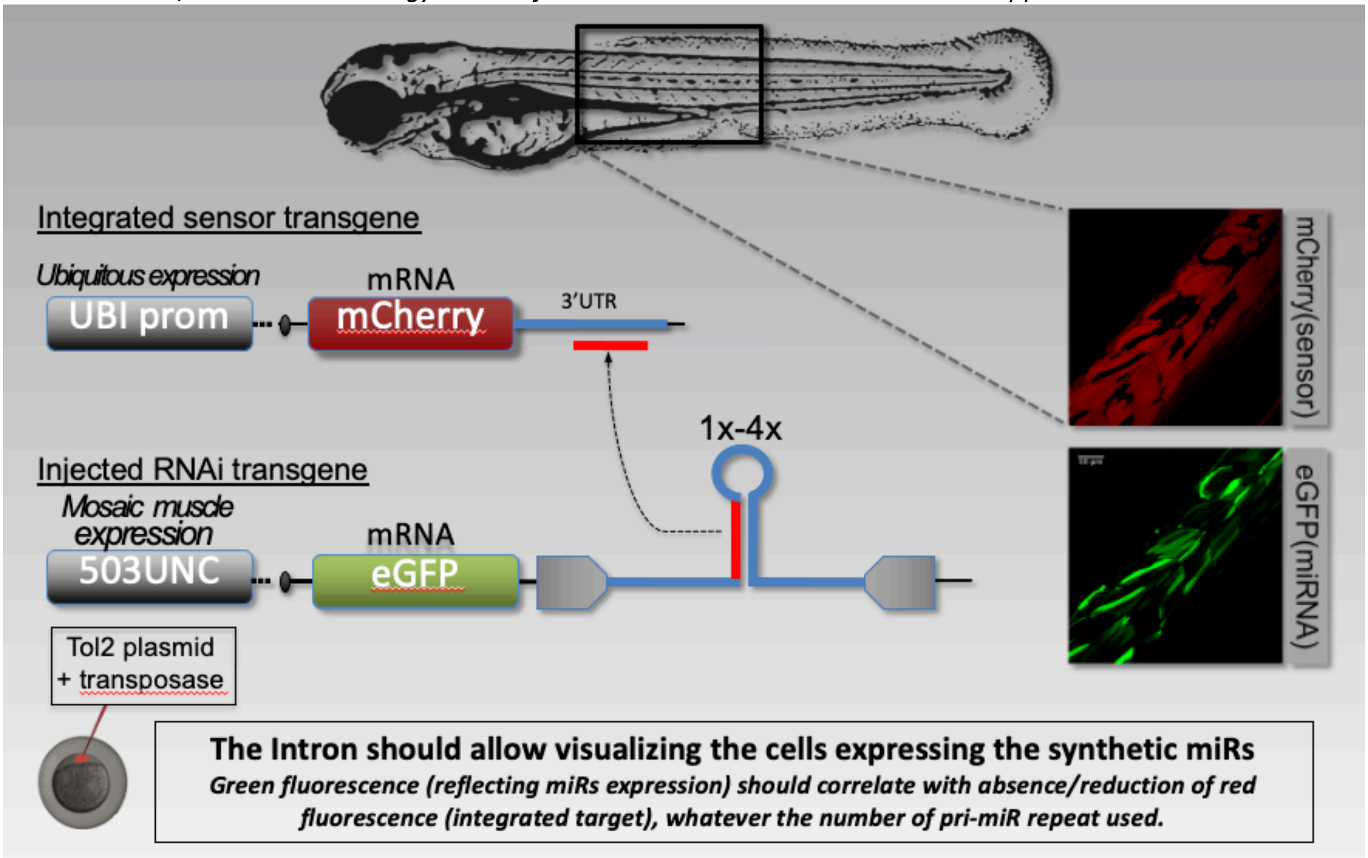


**Figure S03: Schematic representation of the approach used to assess/validate the effect of the intronic sequence.**

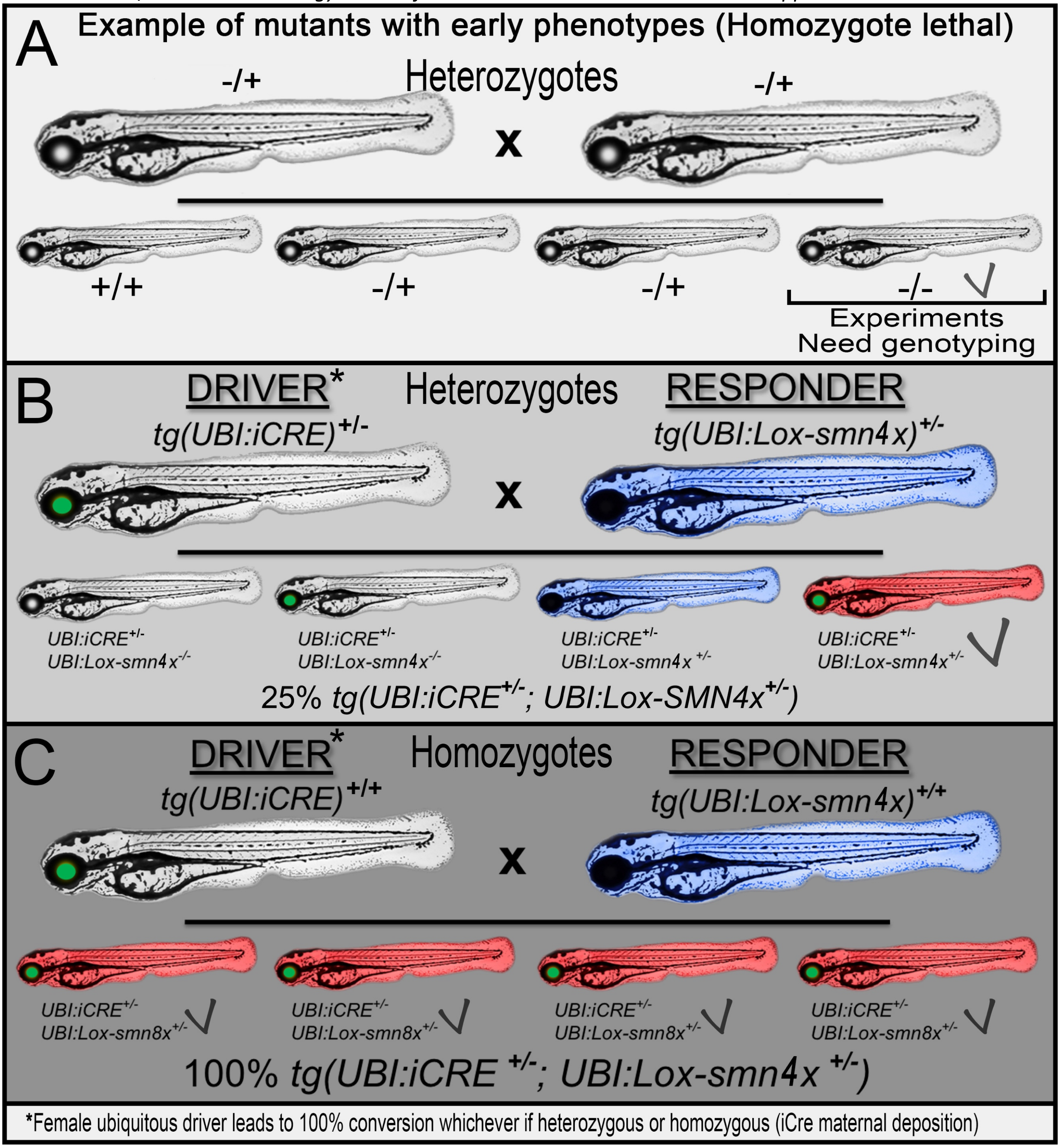


**Figure S04: Illustration presenting crossing required to generate embryos for drug screening or large-scale experiments. A,** Usually, zebrafish disease models of human diseases are suitable for drug screening if they present early and strong phenotypes that can be analysed in multi-well plates. Unfortunately, for models based on mutations, these defects mean that the line can only be maintained in heterozygous state. To generate embryos for the screens, one should incross heterozygous, leading to only 25% of affected homozygous embryos that cannot be identified from their unaffected sibling before the end of the experiments; strongly hampering large-scale experiments. **B-C**, The presented Cre/Lox system can greatly facilitate the generation of affected animals as presented in the schematics. First, the animal could be easily identified before the screening procedure thanks their fluorescent profile. Secondly, and most importantly, this conditional system enables to generate 100% affected animals, strongly easing the experimental procedure. Working with transgenic responder lines leading to strong knockdown while presenting only one insertion is however important for the homogeneity of the screens. Note that female Cre-driver with maternal Cre-expression would lead to 100% conversion even in the absence of driver-transgene transmission.

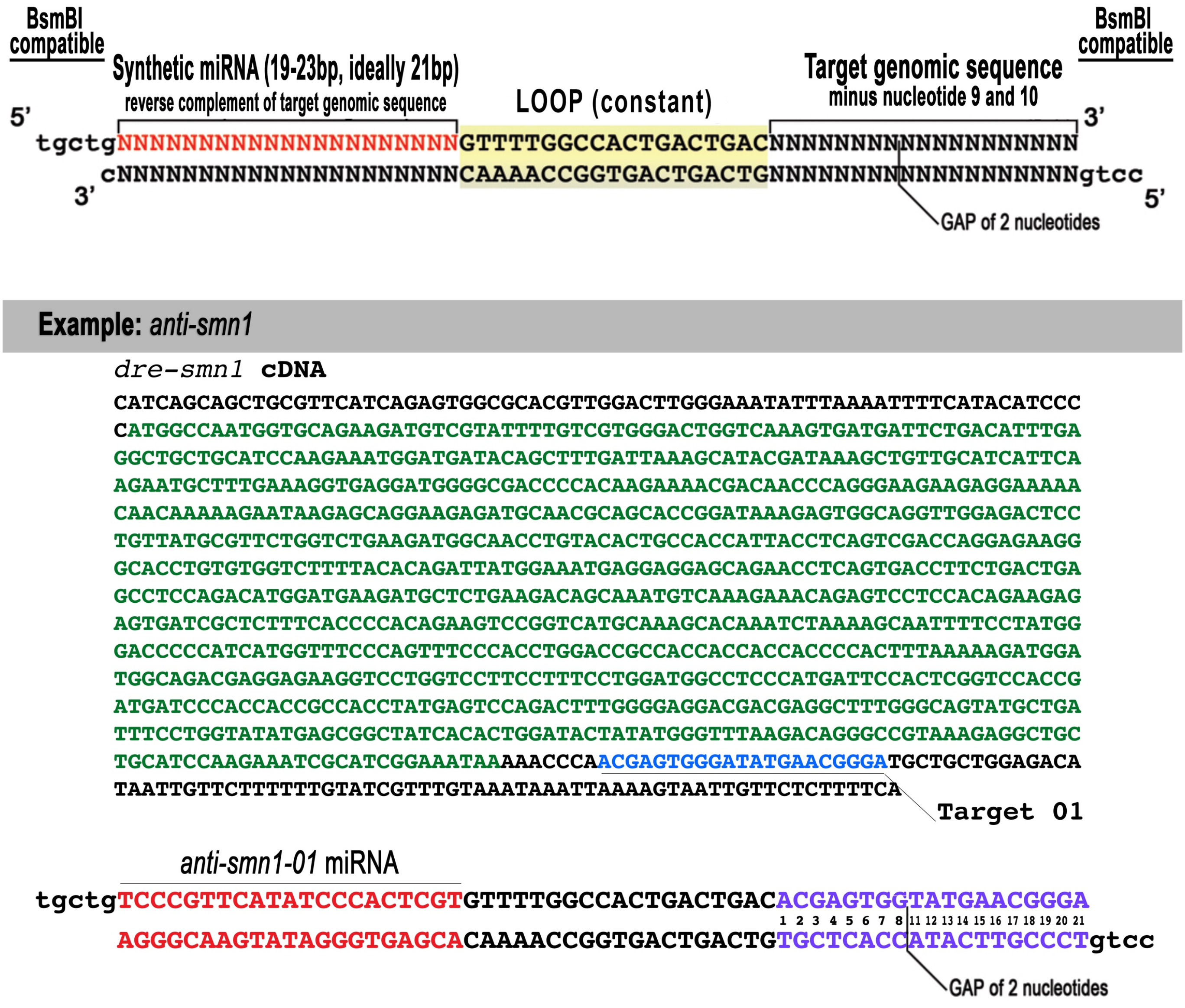


**Figure S05. Synthetic *pri-miRNA* design.** Top and bottom oligo to order for generating (by annealing) *pri-miR* to be inserted into BsmBI-digested p3E-IntronMIR / p3E-IntronRNAi. Sequence in red should be the perfect reverse complement of the DNA target sequence. See the example to target the gene *smn1*.
